# Gene cluster conservation identifies melanin and perylenequinone biosynthesis pathways in multiple plant pathogenic fungi

**DOI:** 10.1101/379305

**Authors:** Malaika K. Ebert, Rebecca E. Spanner, Ronnie de Jonge, David J. Smith, Jason Holthusen, Gary A. Secor, Bart P.H.J. Thomma, Melvin D. Bolton

## Abstract

Perylenequinones are a family of structurally related polyketide fungal toxins with nearly universal toxicity. These photosensitizing compounds absorb light energy which enables them to generate reactive oxygen species that damage host cells. This potent mechanism serves as an effective weapon for plant pathogens in disease establishment. The sugar beet pathogen *Cercospora beticola* secretes the perylenequinone cercosporin during infection. We have shown recently that the cercosporin toxin biosynthesis *(CTB)* gene cluster is present in several other phytopathogenic fungi, prompting the search for biosynthetic gene clusters (BGCs) of structurally similar perylenequinones in other fungi. Here, we report the identification of the elsinochrome and phleichrome BGCs of *Elsinoё fawcettii* and *Cladosporium phlei,* respectively, based on gene cluster conservation with the *CTB* and hypocrellin BGCs. Furthermore, we show that previously reported BGCs for elsinochrome and phleichrome are involved in melanin production. Phylogenetic analysis of the corresponding melanin polyketide synthases (PKSs) and alignment of melanin BGCs revealed high conservation between the established and newly identified *C. beticola, E. fawcettii,* and *C. phlei* melanin BGCs. Mutagenesis of the identified perylenequinone and melanin PKSs in *C. beticola* and *E. fawcettii* coupled with mass spectrometric metabolite analyses confirmed their roles in toxin and melanin production.

**Originality and significance statement:** Genes involved in secondary metabolite (SM) production are often clustered together to form biosynthetic pathways. These pathways frequently have highly conserved keystone enzymes which can complicate allocation of a biosynthetic gene cluster (BGC) to the cognate SM. In our study, we utilized a combination of comparative genomics, phylogenetic analyses and biochemical approaches to reliably identify BGCs for perylenequinone toxins and DHN-melanin in multiple plant pathogenic fungi. Furthermore, we show that earlier studies that aimed to identify these perylenequinone pathways were misdirected and actually reported DHN-melanin biosynthetic pathways. Our study outlines a reliable approach to successfully identify fungal SM pathways.

## Introduction

Fungi produce a plethora of secondary metabolites (SMs) that serve to enhance competitiveness in nature. Functional diversity of these compounds is high, including reported roles in virulence, biotic and abiotic stress protection, and as metal transport agents (Williams et al., 1989; Demain and Fang, 2000; Rohlfs and Churchill, 2011; Stergiopoulos et al., 2012; Keller, 2015). For example, in some occasions SMs are involved in symbiotic relationships where microbial symbionts provide an antibiotic armory against secondary infection to the symbiotically colonized plant in return for nutrients and protection (Rohlfs and Churchill, 2011). A major class of fungal SMs are the polyketides (Keller et al., 2005). For the biosynthesis of fungal aromatic polyketides, non-reducing polyketide synthases (NR-PKSs) play a central role as mediators of the first biosynthetic step (Keller et al., 2005; Crawford and Townsend, 2010; Brakhage, 2013; Gallo et al., 2013). Such PKS genes contain multiple domains that work conjointly, of which the β-ketoacyl synthase (Guedes and Eriksson), acyltransferase (AT), and acyl-carrier protein (ACP) domain are indispensable (Kroken et al., 2003; Keller et al., 2005; Crawford and Townsend, 2010; Gallo et al., 2013). By using the domains iteratively, a PKS generates a metabolite backbone which can be modified by other enzymes to yield the final metabolite (Keller et al., 2005; Bohnert et al., 2010; Crawford and Townsend, 2010). The genes encoding these decorating enzymes are often found in direct proximity to the PKS gene to form a biosynthetic gene cluster (BGC) pathway (Keller and Hohn, 1997; Keller et al., 2005). In addition, BGCs often contain regulatory elements and transporters involved in shuttling the final secondary metabolite from the cell, and in the case of toxic metabolites, genes encoding auto-resistance proteins (Keller, 2015; de Jonge et al., 2018).

A well-studied BGC is the cercosporin toxin biosynthesis *(CTB)* pathway. The *CTB* gene cluster was originally identified in *Cercospora nicotianae,* causal agent of leaf spot disease on tobacco, but is present in almost all *Cercospora* species (Assante et al., 1977; Choquer et al., 2005; de Jonge et al., 2018). The ubiquitous presence of the *CTB* gene cluster in the genus is likely explained by its role as a virulence facilitator (Callahan et al., 1999; Daub and Ehrenshaft, 2000; Choquer et al., 2005). Recently, de Jonge et al. (2018) used comparative genomics to show that the *CTB* gene cluster can also be found in several plant pathogenic fungal species outside the *Cercospora* genus, likely as a result of horizontal transfer of the entire *CTB* gene cluster (Bohnert et al., 2010; Crawford and Townsend, 2010; de Jonge et al., 2018). The majority of assessed species from the genus *Colletotrichum,* a large genus of crop and/or ornamental plant pathogens (Perfect et al., 1999), were shown to harbor full-to partial-length *CTB* gene clusters, of which the post-harvest apple fruit pathogen *Co. fioriniae* was shown to produce cercosporin (de Jonge et al., 2018). The core gene of the *Cercospora CTB* gene cluster is the NR-PKS gene *CTB1* (Newman and Townsend, 2016), which is flanked at both sides by nine genes that putatively encode decorating enzymes (CTB2, CTB3, CTB5, CTB6, CTB7, CTB9, CTB10, CTB11 and CTB12) (de Jonge et al., 2018). Besides those ten genes essential for toxin formation, the cluster also encodes a zinc finger transcription factor (CTB8) for regulation of cluster gene expression, and two major facilitator superfamily (MFS) transporters; CTB4 that is necessary for toxin secretion and the cercosporin facilitator protein (CFP) involved in toxin auto-resistance (Chen et al., 2007; Choquer et al., 2007; de Jonge et al., 2018). Upon activation, all *CTB* pathway enzymes work in a well-orchestrated manner to synthesize the metabolite from backbone formation to secretion of the toxin into the environment whilst providing the fungus with protection against cercosporin.

Cercosporin is a member of the perylenequinone family that, upon photo-activation, displays almost universal toxicity to a wide spectrum of organisms (Zhenjun and Lown, 1990; Daub and Ehrenshaft, 2000; Ahonsi et al., 2005; Guedes and Eriksson, 2007; Daub et al., 2013). Exposure to visible and near-UV light energetically activates perylenequinones to an excited triplet state that reacts with oxygen to form reactive oxygen species (Foote, 1976; Guedes and Eriksson, 2007). This photodynamic activity can be attributed to the 3,10-dihydroxy-4,9-perylenequinone chromophore backbone that is shared among perylenequinones (Hudson et al., 1997). Structural differences between perylenequinone family members are mostly due to divergent side chains attached to the mutual backbone structure (Daub et al., 2005) (Fig. 1). For example, the methylenedioxy bridge is a unique feature of cercosporin and is absent in other perylenequinones such as hypocrellin, elsinochrome and phleichrome (Fig. 1) (Weiss et al., 1987; de Jonge et al., 2018).

**Figure 1.**
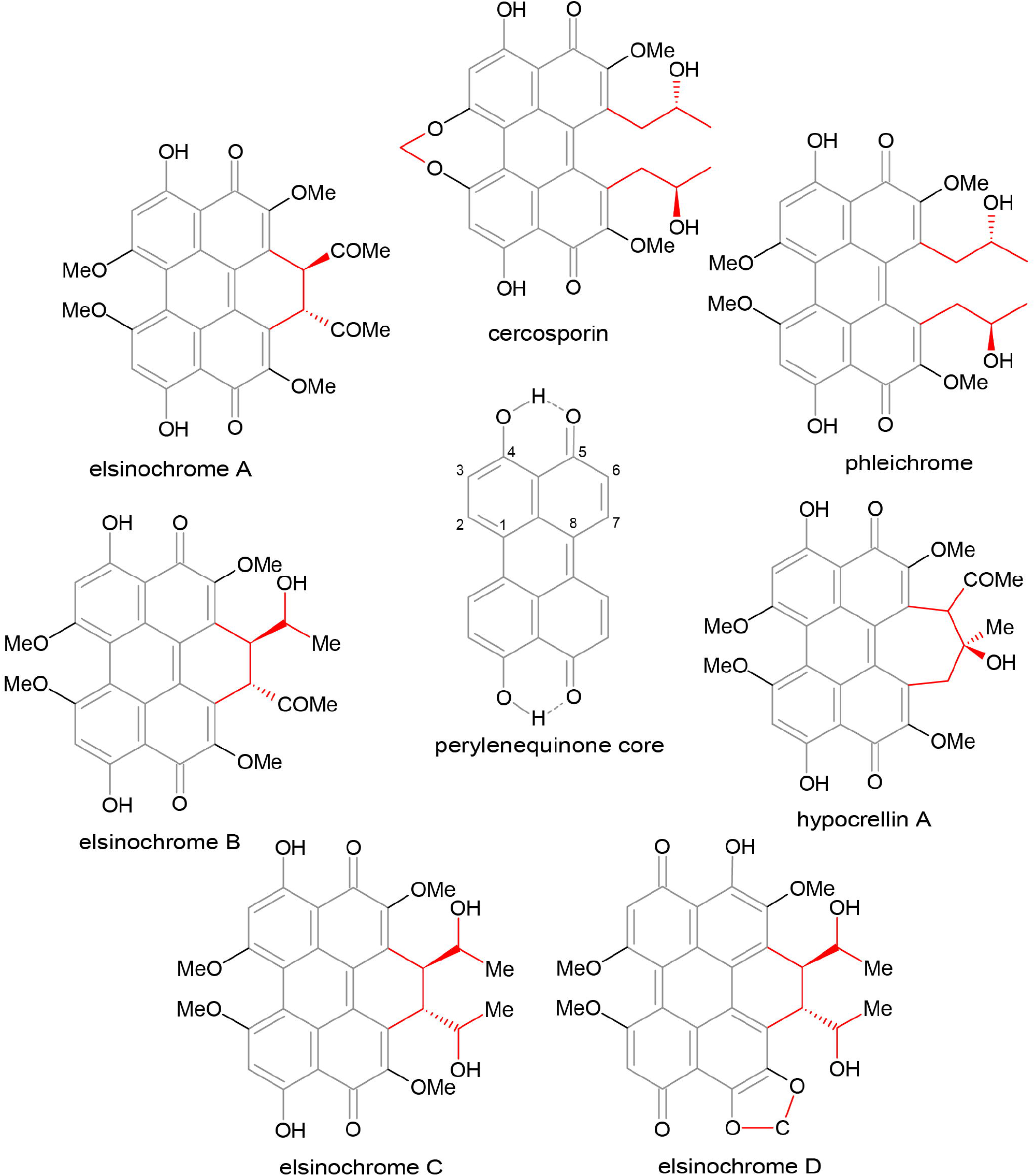
Structures of related perylenequinones. Cercosporin synthesized by *Cercospora* spp., phleichrome by *C. phlei* and elsinochromes A, B, C, and D produced by *E. fawcettii* are structurally related as they share a common perylenequinones backbone (center; indicated in the molecule structure in gray). Structural differences between the molecules are mostly due to various side chains (indicated in red). Differences between perylenequinones are observed at positions 2, 2’and 7, 7’.

Previous studies have implicated PKS genes in the production of perylenequinones in other plant pathogenic fungi. For example, transcriptome analysis and a CRISPR-Cas9 gene editing approach in the bamboo pathogen *Shiraia bambusicola* gave compelling evidence that *SbaPKS* encodes the PKS orchestrating hypocrellin biosynthesis (Deng et al., 2017, Zhao et al., 2016). Similarly, targeted disruption of *EfPKSl* in the citrus scab pathogen *Elsinoë fawcettii* appeared to abrogate elsinochrome production (Liao and Chung, 2008). Likewise, Cppks1 was suggested to be responsible for PKS activity for phleichrome production in the purple eyespot pathogen *Cladosporium phlei* (So et al., 2015). However, we have previously used KS domain phylogeny to associate PKS genes with the final perylenequinone product (de Jonge et al., 2018). During the course of these analyses, we identified PKS genes for *E. fawcettii* and *C. phlei* that were not previously attributed to these perylenequinones, which prompted us to re-evaluate the findings of Liao and Chung (2008) and So et al. (2015). Interestingly, Liao and Chung (2008) also carried out phylogenetic analysis of EfPKS1 with other PKSs which indicated that EfPKS1 clustered closely to a diverse set of fungal non-reducing PKSs involved in biosynthesis of the toxins cercosporin, aflatoxin and sirodesmin, but also with PKSs involved in pigment production such as dihydroxynaphthalene (DHN)-melanin. Melanin is an integral component of the cell wall that has proposed functions in protection from environmental factors, appressorial penetration of host plants and pathogenesis (Wheeler and Bell, 1988; Langfelder et al., 2003; Liu and Nizet, 2009). In *Mycosphaerella fijiensis,* research suggested that secreted fungal DHN-melanin acts as a virulence factor through the photogeneration of singlet molecular oxygen in a similar manner to the perylenequinones (Beltrán-García et al., 2014). DHN-melanin biosynthesis has been characterized extensively in many fungi, including *Magnaporthe oryzae, Colletotrichum lagenarium, Alternaria alternata, Botrytis cinerea, Verticillium dahliae* and *Aspergillus* spp.. In the rice blast fungus *M. oryzae* for instance, DHN-melanin production is known to be mediated by a four-gene cluster which is regulated in hyphae by the transcription factor Pig1 (Fig. 2) (Thompson et al., 2000; Tsuji et al., 2000; Talbot, 2003; Oh et al., 2008). However, fungal DHN-melanin pathways may vary in the biosynthesis of the first common intermediate 1,3,6,8-tetrahydroxynaphthalene (1,3,6,8-THN or T4HN). For example, the PKS ALB1 (for “albino 1”) is responsible for the first biosynthetic step in *Aspergillus fumigatus,* resulting in the biosynthesis of the heptaketide naphthopyrone YWA1, which is subsequently hydrolyzed by Ayg1 to produce T4HN (Fujii et al., 2004; Pihet et al., 2009). Two alternative routes can be found in the necrotrophic gray mold fungus *B. cinerea.* In this case, the PKSs Bcpks12 and Bcpks13 synthesize different precursors for the joint DHN-melanin pathway (Schumacher, 2016). While Bcpks12 produces the pentaketide T4HN directly, Bcpks13 synthesizes the hexaketide 2-acetyl-1,3,6,8-tetrahydroxynaphthalene (AT4HN) that is subsequently converted to yield T4HN (Schumacher, 2016). In either case, the resulting T4HN will serve as substrate for a hydroxynaphthalene (HN) reductase leading to scytalone formation. In the next step, scytalone will be dehydrated by a scytalone dehydratase resulting in the formation of 1,3,8-trihydroxynapthalene (1,3,8-THN or T3HN). Subsequent reduction by a HN reductase yields vermelone which is subsequently dehydrated to form 1,8-DHN; an immediate precursor of melanin (Tsai et al., 1999; Thompson et al., 2000; Tsuji et al., 2000; Talbot, 2003).

**Figure 2.**
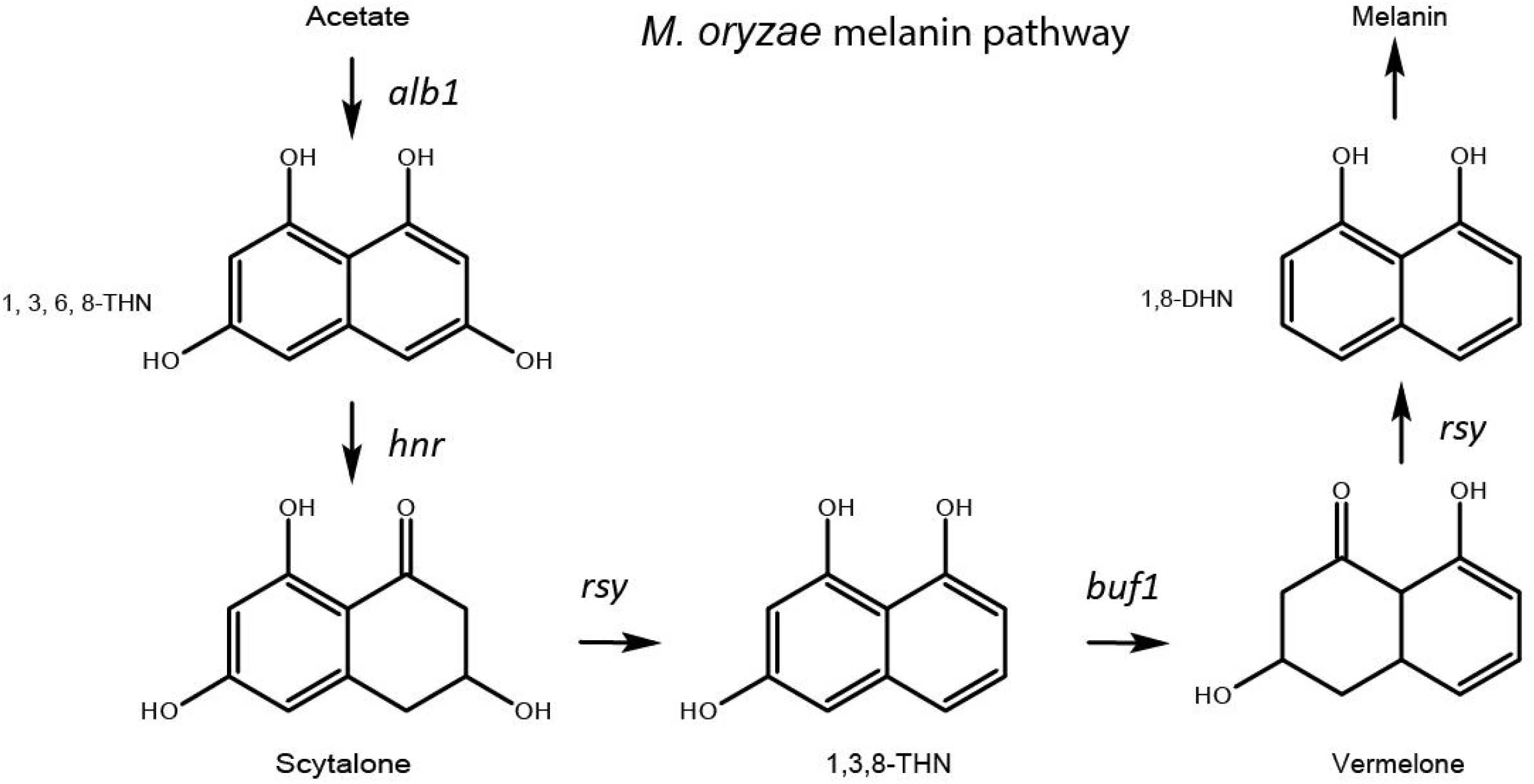
Schematic DHN-melanin biosynthesis pathway of *M. oryzae*. In the first biosynthetic step, the PKS ALB1 forms 1,3,6,8-tetrahydroxynaphthalene (1,3,6,8-THN or T4HN) by ketide cyclization. Reduction by the tetrahydroxynaphthalene reductase 4HNR results in the formation of scytalone which will be dehydrated by RYS1, a scytalone dehydratase, to yield trihydroxynaphthalene (T3HN). The T3HN reductase BUF1 subsequently reduces T3HN to vermelone followed by a dehydration step mediated by RYS1 to form dihydroxynaphthalene (2HN), the immediate precursor of melanin.

In this manuscript, we show that the gene clusters housing *Cppks1* and *EfPKS1* have high similarity to established gene clusters involved in DHN-melanin biosynthesis and have only limited similarity to the perylenequinone biosynthesis clusters to which they were previously attributed. Due to its detailed characterization, the established *M. oryzae* melanin cluster was used as reference in our alignments of putative DHN-melanin gene clusters of *C. beticola, E. fawcettii, C. phlei,* and *S. bambusicola* to illustrate the high level of homology among the DHN-melanin BGCs. Consequently, we also sought to establish the BGCs involved in production of elsinochrome in *E. fawcettii* and phleichrome in *C. phlei,* and included targeted gene replacement of both perylenequinone and melanin PKS genes in *E. fawcettii* and *C. beticola* to provide proof for their involvement in toxin and DHN-melanin production.

## Results

### E. fawcettii *and* C. phlei *genomics*

Nuclear and mitochondrial DNA of *E. fawcettii* strain CBS 139.25 and *C. phlei* strain CBS 358.69 were sequenced to approximately 138-fold and 110-fold coverage, respectively, on the Illumina HiSeq 4000 platform (paired-end, 100-bp reads). Raw reads were processed and assembled by SPAdes (version 3.9.0) yielding draft genome assemblies of 25.3 Mb on 398 scaffolds for *E. fawcettii* and 31.9 Mb on 794 scaffolds for *C. phlei.* The respective scaffold N50 values and L50 numbers for these assemblies are 13 and 676 Kb, and 44 and 238 Kb. Following genome assembly, we used Augustus (version 3.2.1) with default settings (Stanke et al., 2008) and the previously devised *C. beticola* training parameters (de Jonge et al., 2018) to predict 9,519 (mean length 1,675 bp and ~2.5 exons/gene) and 11,316 (mean length 1,624 bp and ~2.3 exons/gene) protein-coding genes for *E. fawcettii* and *C. phlei* respectively. Finally, protein function as well as putative localization was predicted by Interpro (Finn et al., 2017) scanning and yielded annotations for 9,253 out of 9,519 *E. fawcettii* proteins and 10,870 out of 11,316 *C. phlei* proteins. Considering only hits to Pfam, SMART, CDD or SUPERFAMILY databases, 7,450 (78%) and 8,479 (75%) genes were annotated for *E. fawcettii* and *C. phlei,* respectively.

### PKS genealogy and prediction of function

To study the level of conservation of PKSs and associated pathways involved in the biosynthesis of different perylenequinones, we mined the genomes of both perylenequinone producers and non-producers for non-reducing PKSs (Suppl. Table 1). Subsequently, the phylogenetic relationships between these PKSs and those of previously characterized PKSs from selected species as found in Collemare et al. (2014) (Suppl. Table 1) were determined by aligning the highly conserved β-ketoacyl synthase (Guedes and Eriksson) domains of each PKS (Fig. 3). This genealogy revealed distinctive clade formation where PKSs with confirmed involvement in biosynthesis of structurally similar metabolites were observed to cluster. The clades were categorized as perylenequinone, aflatoxin, anthraquinone, or DHN-melanin biosynthesis depending on the function of confirmed PKSs they harbored (Fig. 3). Interestingly, the PKSs EfPKS1 (Liao and Chung, 2008)/ [ELSFAW09157-RA (this study)] from *E. fawcettii* and Cppks1 (So et al., 2015)/[CLAPH08786-RA (this study)] from *C. phlei* that were previously implicated in perylenequinone biosynthesis did not cluster phylogenetically with the established perylenequinone cercosporin PKSs CbCTB1 and CnCTB1 of *C. beticola* and *C. nicotianae,* respectively. Instead, EfPKS1 and Cppks1 formed a clade with confirmed melanin PKSs, including Bcpks12 and Bcpks13 of the gray mold fungus *B. cinerea* (Schumacher, 2016), Wdpks1 of the zoopathogenic black yeast *Wangiella (Exophiala) dermatitidis* (Feng et al., 2001), GIPKS1 of the filamentous fungus *Glarea lozoyensis* (Zhang et al., 2003), NodPKS1 of an endophytic *Nodulisporium* strain (Fulton et al., 1999), COGPKS1 of the cucumber anthracnose causal agent *Co. lagenarium* (Fujii et al., 1999), the predicted *C. beticola* melanin biosynthesis PKS *CbPKS1* (CBET3_09638) and the *S. bambusicola* melanin PKS SHIR08477. The finding that *E. fawcettii* ELSFAW09157-RA, *C. phlei* CLAPHL08786-RA, *S. bambusicola* SHIR08477, and *CbPKS1* reside in a cluster (Fig. 3) with extensive collinearity to established DHN-melanin clusters (Fig. 4) suggests a role in melanin production and hints that *EfPKS1* and *Cppks1* were previously misannotated as perylenequinone biosynthesis genes (Liao and Chung, 2008; So et al., 2015).

**Figure 3.**
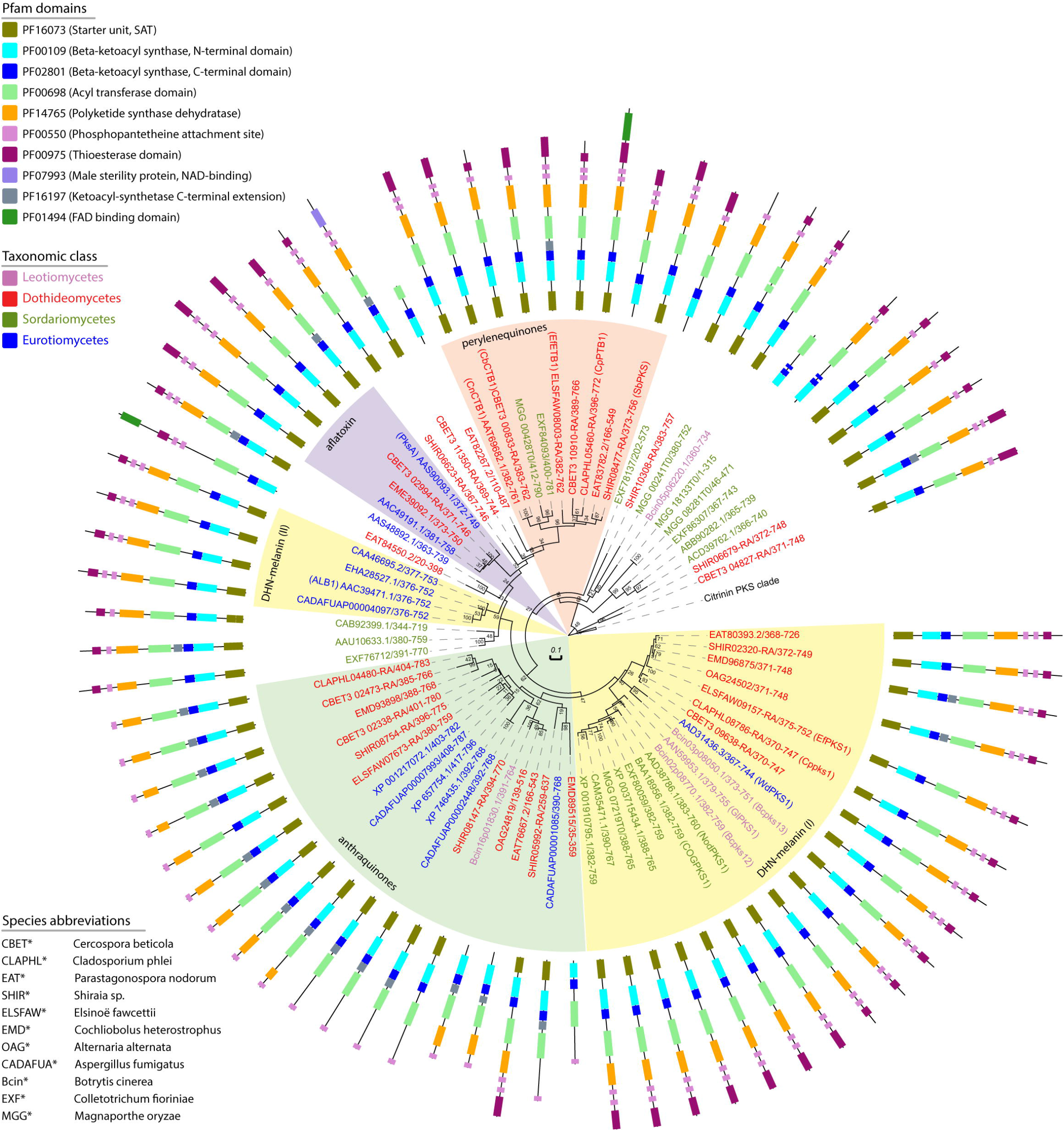
Phylogeny of PKS genes of related Ascomycetes revealing distinct DHN-melanin and perylenequinone subclades. Circular maximum likelihood phylogenetic tree illustrating the phylogenetic relationship of all predicted non-reducing polyketide synthase (PKS) from the selected species set (Supl. Table 1) plus those derived from the set of PKSs used by Collemare et al. (2014). The tree was constructed by maximum-likelihood analysis of aligned full-length β-ketoacyl synthase domains. The outside ring indicates domain architecture of each PKS determined by Pfam domain annotation. Protein accessions are colored depending on the taxonomic class of the producing species. The corresponding species identity for each protein can be found in the bottom left corner. Established biosynthetic end products for a subset of the listed PKSs is indicated by the background color, highlighting two DHN-melanin sub-groups, naphthoquinones, anthraquinones, perylenequinones, aflatoxin-like compounds and resorcylic acid lactones.

**Figure 4.**
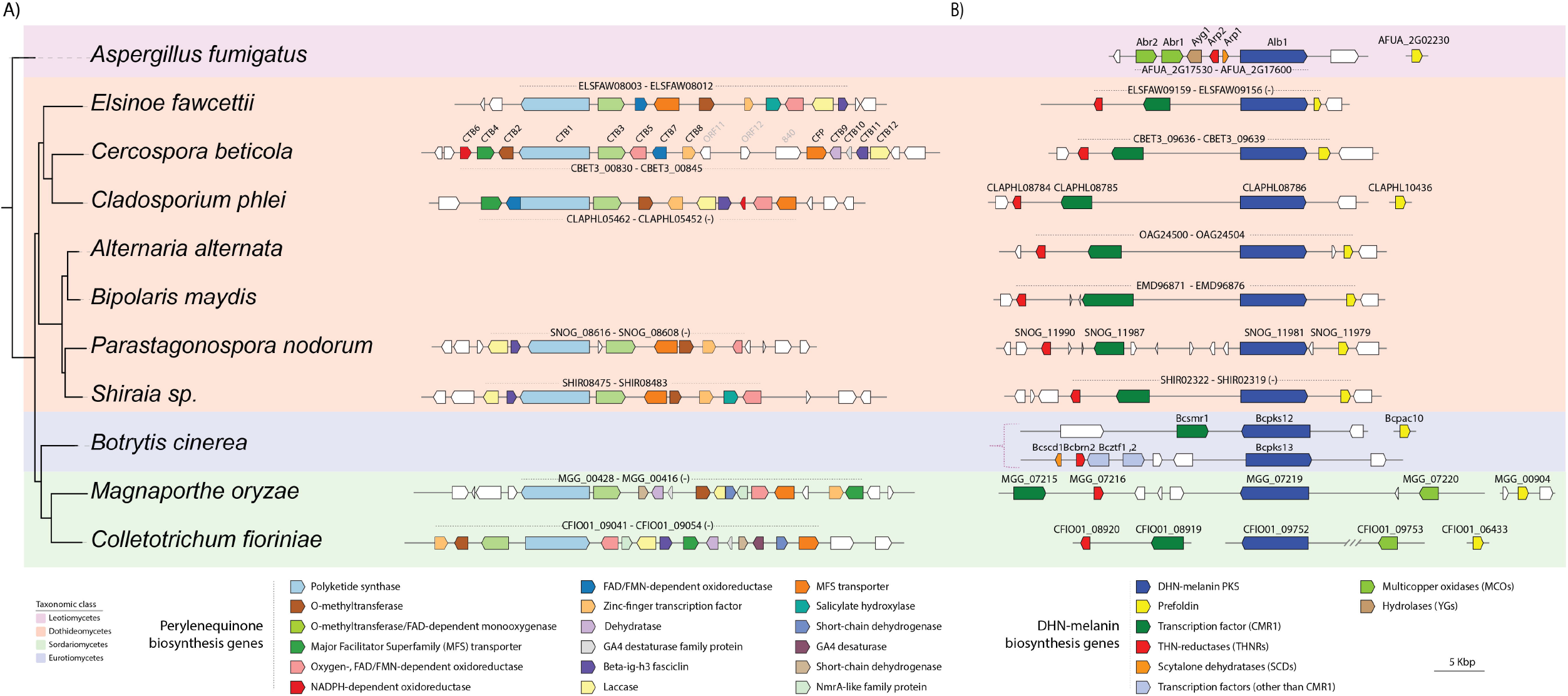
Synteny and rearrangements of conserved perylenequinone and DHN-melanin BGCs. Phylogenetic tree of Ascomycetes used in this study based on mash protein-level kmer hash overlaps. Alignments were based on MultiGeneBlast with a selected set of genomes and an input BGC as query. Alignment of established and putative perylenequinone BGCs of *E. fawcettii, C. beticola, C. phlei, P. nodorum, S. bambusicola, M. oryzae,* and *Co. fioriniae* (A). For the DHN-melanin BGC alignment, putative and established DHN-melanin BGCs of *E. fawcettii, C. beticola, C. phlei, P. nodorum, S. bambusicola, M. oryzae, Co. fioriniae, A. fumigatus, A. alternata, B. maydis (C. heterostrophus),* and *B. cinerea* were aligned (B). For all species, the indicated identifiers are transcript IDs and the corresponding sequences can be retrieved from Ensemble Fungi and/or NCBI GenBank. *CTB* orthologs are colored relative to the *C. beticola CTB* cluster genes while DHN-melanin BGC genes are color coded relative to *M. oryzae* DHN-melanin. Color key and annotated functions are explained in the legend below the cluster graphics.

**Figure 5.**
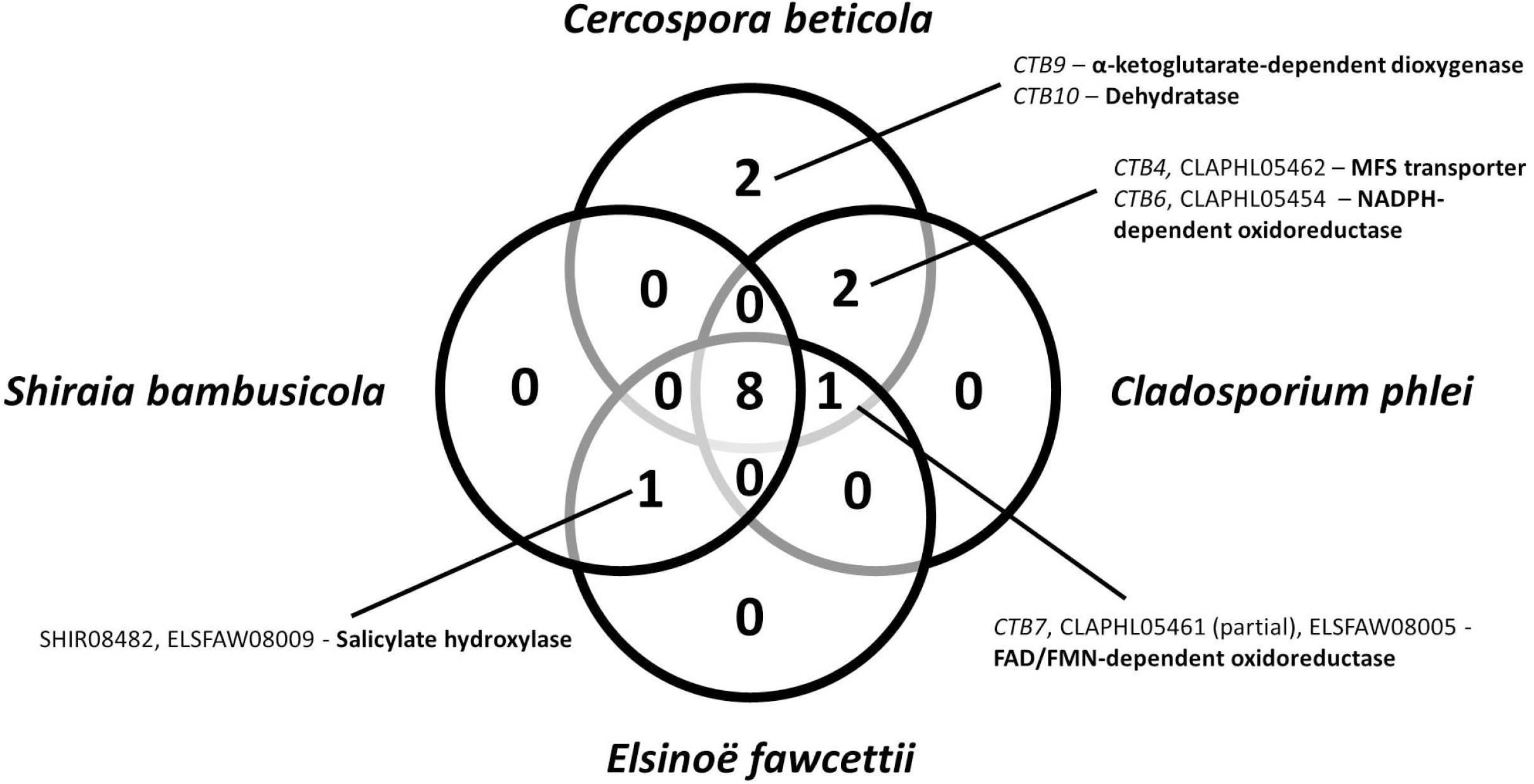
Conserved and unique genes in the confirmed or predicted perylenequinone BGCs of *C. beticola, E. fawcettii, C. phlei,* and *S. bambusicola.* Venn diagram highlights the number of shared BGC genes of the cercosporin, elsinochrome, phleichrome, and hypocrellin pathways.

The cercosporin PKSs in *C. beticola* (CbCTB1), *C. nicotianae* (CnCTB1), and *Co. fioriniae* (EXF84093) form a perylenequinone clade with the previously confirmed hypocrellin PKS (SbaPKS) (Zhao et al., 2016; Deng et al., 2017), ELSFAW08003 from *E. fawcettii,* CLAPHL05460 from *C. phlei* as well as with the putative perylenequinone PKSs in *P. nodorum* (EAT83782.2), *M. oryzae* (MGG_00428), and the *C. beticola* CbCTB1 paralog CBET3_10910 (Fig. 3). As phylogenetic conservation can be an indication of related metabolite production (de Jonge et al., 2018), this clustering suggests that PKSs of this clade are involved in biosynthesis of the perylenequinones. Therefore, we suggest renaming *ELSFAW08003* to *EfETB1* for <u>e</u>lsinochrome <u>t</u>oxin <u>b</u>iosynthesis gene 1, and *C. phlei CLAPHL05460* to *CpPTB1* for <u>p</u>hleichrome <u>t</u>oxin <u>b</u>iosynthesis <u>g</u>ene 1.

### Perylenequinone and DHN-melanin biosynthesis gene cluster alignments

While PKS genes are indispensable for polyketide formation, it is the full complement of genes in a BGC that is responsible for the biosynthesis of the end product. Therefore, synteny of the predicted BGCs of orthologous PKS genes was assessed. Using the established *C. beticola CTB* gene cluster and *S. bambusicola* hypocrellin gene cluster as references, putative perylenequinone orthologous gene clusters in *E. fawcettii, C. phlei, P. nodorum, M. oryzae,* and *Co. fioriniae* were aligned (Fig. 4A). Although there is evidence for gene loss and gain between the perylenequinone BGC alignments, multiple core genes are shared between cercosporin, hypocrellin, and the predicted BGCs for elsinochrome and phleichrome (Fig. 4A). Overall, eight genes are shared between the cercosporin, hypocrellin and predicted elsinochrome and phleichrome BGCs (Figs. 4A, 5). When compared to these perylenequinone pathways, the *CTB* gene cluster has two additional genes; a putative α-ketoglutarate-dependent dioxygenase *(CTB9)* and a candidate dehydratase *(CTB10)* that have been shown to be involved in the formation of the methylenedioxy bridge (de Jonge et al., 2018). The predicted *C. phlei* phleichrome BGC contains all orthologous *C. beticola CTB* genes except for the above-mentioned *CTB9* and *CTB10,* in agreement with the lack of the methylenedioxy bridge in phleichrome. Likewise, the predicted *E. fawcettii* elsinochrome BGC lacks *CTB9* and *CTB10* as well as the cercosporin MFS transporter *(CTB4)* and the NADPH-dependent oxidoreductase *(CTB6).* Interestingly, the *E. fawcettii* BGC contains *ELSFAW08009,* which only has an ortholog in the hypocrellin gene cluster *(SHIR08482)* and in no other of the aligned BGCs (Fig. 4A). *ELSFAW08009* and *SHIR08482* are annotated as a putative salicylate hydroxylase based on sequence similarity to the conserved protein domain family TIGR03219 (E-value 2.98e-18), members of which are salicylate 1-monoxygenases. Besides sharing this gene with the elsinochrome pathway and lacking orthologs to *CTB9* and *CTB10,* the hypocrellin cluster also lacks CTB homologs *CTB4, CTB6,* and *CTB7* compared to the cercosporin pathway (Fig. 4A).

Similarly, predicted DHN-melanin clusters of *C. beticola, C. phlei, E. fawcettii, S. bambusicola sp. slfl4,* and *Co. fioriniae* were aligned to the established DHN-melanin cluster of *M. oryzae, A. fumigatus, A. alternata, Bipolaris maydis (Cochliobolus heterostrophus),* and both alternative clusters of *B. cinerea* (Figure 4B). All BGCs share homologous PKS genes, a THN-reductase, and a prefoldin-encoding gene. Prefoldins are frequently associated with DHN-melanin BGCs, but a functional role in DHN-melanin biosynthesis has not been established to date. Furthermore, the putative melanin clusters of *C. beticola, C. phlei, E. fawcettii, S. bambusicola sp. slf14,* and *Co. fioriniae* contain a transcription factor with homology to *M. oryzae* Pig1 and Co. *lagenarium* CMR1, which are often observed in other established melanin clusters (Tsuji et al., 2000).

### Targeted replacement and characterization of perylenequinone and melanin PKS genes

The predicted perylenequinone and melanin PKS genes for *C. beticola (CbCTB1* and *CbPKS1,* respectively) and *E. fawcettii (EfETB1* and *EfPKS1,* respectively) were targeted for split marker gene replacement. At least two unique site-directed transformants were assessed for involvement in metabolite production. The wild type and all knockout mutant strains were grown under conditions to induce perylenequinone production. The presence or absence of cercosporin (C. *beticola)* and elsinochrome *(E. fawcettii)* in culture extracts was determined via UPLC-MS (Fig. 6A and B). For both fungal species, perylenequinone production was abrogated in the perylenequinone PKS mutants *(ΔCbCTB1* and *ΔEfETB1* mutants for *C. beticola* and *E. fawcettii,* respectively) but not in the melanin PKS mutants (Fig. 6A and B). There were no obvious differences in growth rate for either of the *C. beticola* or *E. fawcettii* mutants versus the corresponding wild type strains. Additionally, *ΔCbPKS1* and *ΔEfPKS1* melanin mutants had a pale buff color as opposed to the dark grey pigmentation observed in wild type and perylenequinone-deficient mutant *(ΔCbCTB1* and *ΔEfETB1* for *C. beticola* and *E. fawcettii,* respectively) strains (Figure 7A and B). The amount of melanin present in the *C. beticola* cultures was determined spectrophotometrically, showing that the *ΔCbPKS1* mutant had a significantly lower melanin content than the wild type (P < 0.05) (Fig. 7C). A quantitative evaluation of melanin content of *E. fawcettii* wild type and mutant cultures is yet to be reported as earlier extractions did not yield sufficient quantity for comparison.

**Figure 6.**
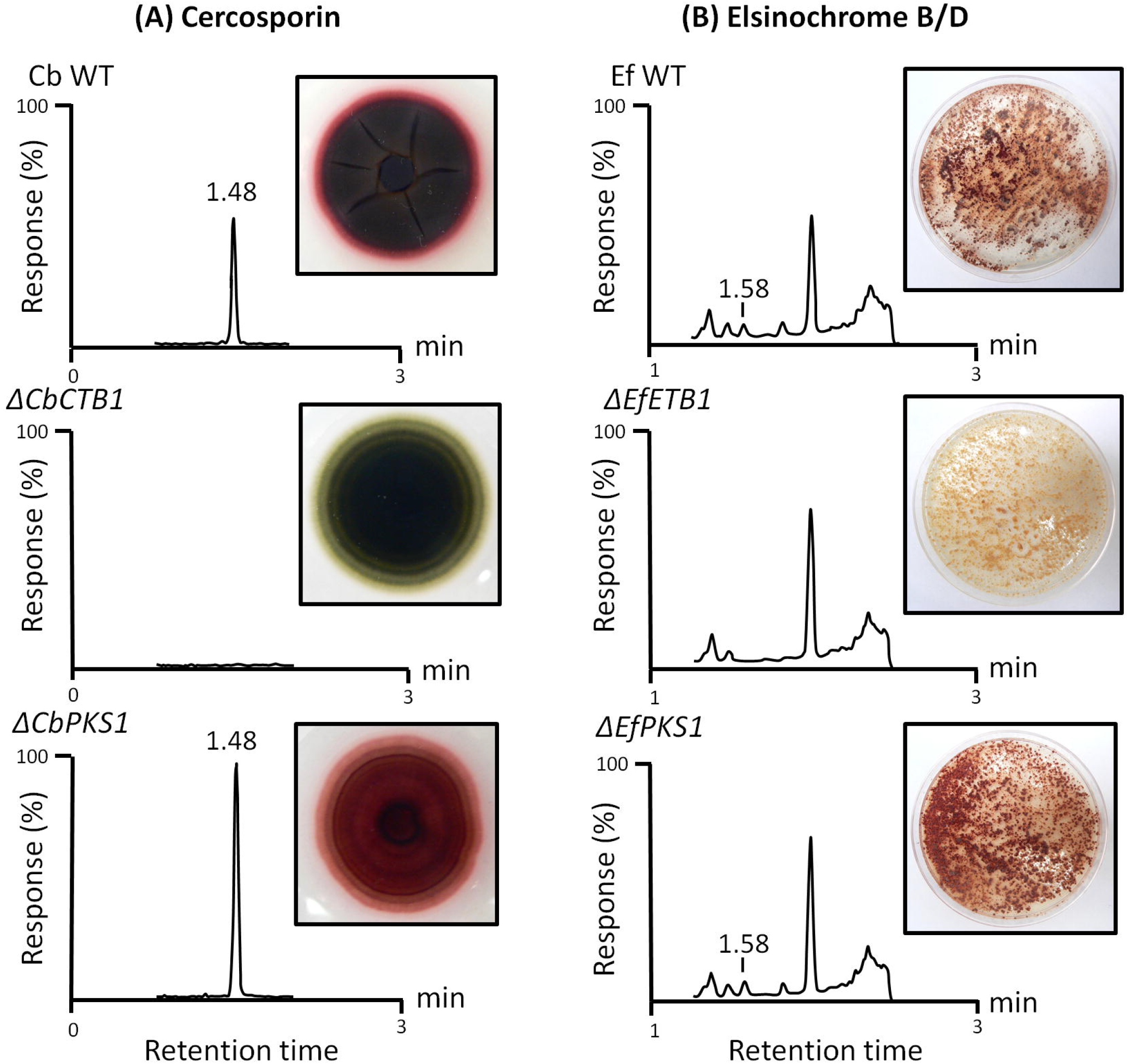
Perylenequinone toxin detection in *C. beticola* and *E. fawcettii* perylenequinone and melanin PKS mutants compared to wild type strains after growth under perylenequinone-inducing conditions. Representative UPLC mass-selective detection of cercosporin (column A) and elsinochrome B/D (column B) are shown for each fungal strain (minimum of 2 plate extracts per strain). Cercosporin (column A) was present in *C. beticola* wild type and the *CbΔPKS1* mutants at a (retention time 1.48 min) but not in the *CbΔCTB1* mutants (the cercosporin standard produced a mass-selective chromatogram with an identical retention time; data not shown). An elsinochrome B/D peak (column B) was present in wild type *E. fawcettii* and *EfΔPKS1* strains, retention time 1.58 min, and was undetectable in *EfΔETB1* mutants (no chemical-grade standard available).

**Figure 7.**
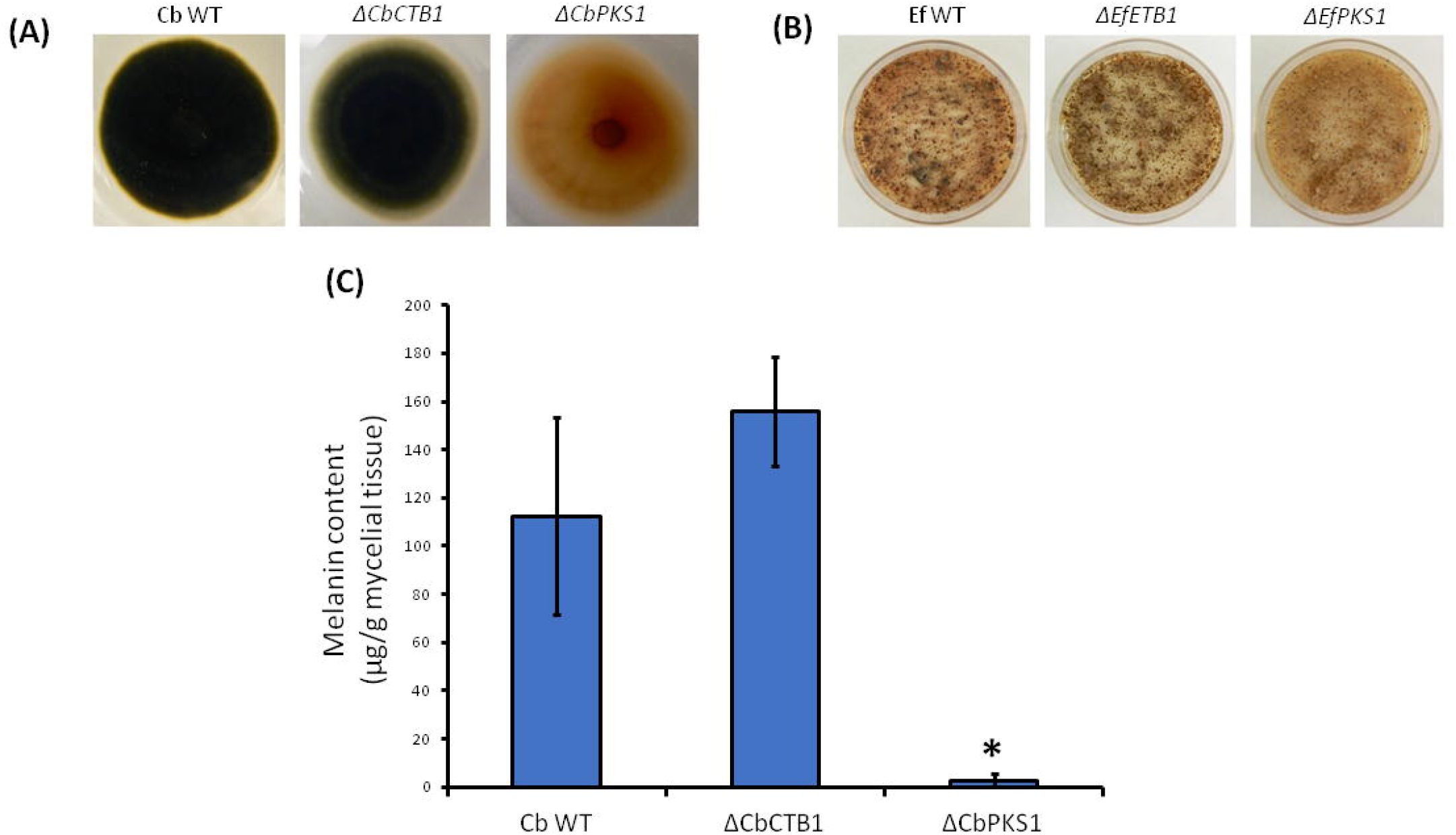
Melanin production in *C. beticola* perylenequinone and melanin PKS mutants compared to wild type. Photos of *C. beticola* strains grown on V8 agar in conditions conducive to sporulation (described in methods) (A). Photos of *E. fawcettii* grown on Fries agar in conditions conducive to melanin production (B). The mean melanin content of three individual fungal cultures (μg melanin/g of mycelial tissue ± standard error) in (C) *C. beticola* wild type (WT), melanin mutants *(CbΔPKS1),* and cercosporin mutants *(CbΔCTB1).* Significant differences (P< 0.05) indicated by *.

## Discussion

Phylogenetic analysis based on PKS KS domain conservation can help to predict SM structure and gene evolution (Keller et al., 2005; Gallo et al., 2013). In this study, we used KS domain sequence alignments and phylogenetic analysis of selected plant pathogenic fungi to separate PKSs into distinct clades. One of the clades hosted PKS genes involved with perylenequinone biosynthesis including *CbCTB1,* the well-studied *Cercospora beticola* PKS essential for cercosporin biosynthesis, and the PKS gene of the hypocrellin pathway in *S. bambusicola sp. slf14.* We also observed clustering of PKS genes involved in DHN-melanin formation such as *Bcpks12* and *Bcpks13* of *B. cinerea* and *COGPKS1* of Co. *lagenarium* (Fig. 3). As previously reported (Liao and Chung, 2008), phylogenetic analyses of KS and AT domain sequences indicated a closer relationship of EfPKS1 to melanin PKSs than to perylenequinone PKSs. Furthermore, high similarity of the full length amino acid sequence to the annotated EfPKS1 led So et al. (2015) to hypothesize that Cppks1 was involved in phleichrome production. Our KS domain alignment confirms the phylogenetic analysis by Liao and Chung (2008) where EfPKS1 and Cppks1 form a cluster with established DHN melanin biosynthesis PKSs of other Ascomycetes (Fig. 3). Consequently, we used comparisons to well-characterized melanin BGCs in various Ascomycetes to show that PKS genes belonging to the DHN-melanin clade are putatively involved with melanin biosynthesis in *C. beticola, E. fawcettii, C. phlei,* and *S. bambusicola sp. slf14* (Fig. 4B). Besides PKS phylogeny, whole-cluster homology of predicted cognate clusters to various well established DHN-melanin clusters strengthened our hypothesis that *CbPKS1, EfPKS1,* and *Cppks1* are involved with melanin production.

To gain further support, we generated *PKS* mutants in our candidate melanin biosynthesis PKS genes in *C. beticola and E. fawcettii.* As predicted, the melanin null mutants Δ*EfPKS1* and Δ*CbPKS1* displayed pale phenotypes characteristic to previously described melanin-deficient mutant strains (Chumley and Valent, 1990) (Fig. 7A and B) and quantitative determination of melanin in Δ*CbPKS1* of *C. beticola* indicated reduced melanin content of the culture compared to wild type and *ΔCbCTB1* knockout mutant lines (Fig. 7C).

Interestingly, the *C. beticola ΔCbPKS1* and *E. fawcettii ΔEfPKS1* mutants were still able to produce cercosporin and elsinochrome, respectively (Fig. 6 A and B) unlike the observation by Liao and Chung (2008) where *ΔEfPKS1* mutants in *E. fawcettii* were reported to be elsinochrome-deficient. Therefore, we suspect that the phenotype they observed may be due to insufficient perylenequinone-inducing conditions. Previous functional analysis of the *EfPKS1* cluster raised questions when overexpression of predicted cluster genes under different conditions did not correlate with elsinochrome production (Chung and Liao, 2008) and complementation of the transcription factor-like gene *EfTSF1* null mutant (with *EfTSF1)* was unable to restore elsinochrome production to wild type level (Liao and Chung, 2008). Furthermore, Chung and Liao (2008) stated that positive *ΔTSF1* transformants were selected based on elsinochrome-deficient phenotype, potentially dismissing any true site-directed mutants that were still able to produce elsinochrome. Therefore, it could be that their elsinochrome-deficient phenotype was evoked by off target tDNA insertion. The reduction in virulence observed for their *ΔEfPKS1* mutant is not surprising as melanin has been reported to be a virulence factor for many filamentous fungi (Langfelder et al (2003); Liu and Nizet (2009); Wheeler and Bell, 1988). Besides contribution to fungal virulence, melanin has also been reported to play an important role in protection against environmental stresses. Recently, studies of the causal agent of Septoria tritici blotch on wheat, *Zymoseptoria tritici,* have indicated a correlation between fungicide resistance and melanization level of the producing fungus which led to the identification of the putative *Z. tritici* melanin PKS (Lendenmann et al., 2014; Lendenmann et al., 2015). Similarly, *CbPKS1* and *CBET3_09636,* encoding a predicted tetrahydroxynaphthalene (T4HN) reductase (now renamed to *Cb4HNR* as it is homologous to *4HNR* of *M. oryzae),* that we propose to belong to the melanin BGC have been recently reported to be more highly expressed in fungicide-resistant *C. beticola* strains compared to fungicide-sensitive strains (Bolton et al., 2016). Consequently, we propose that melanin production in *C. beticola* is mediated by *CbPKS1* which forms T4HN in the first biosynthetic step. Subsequently, T4HN will serve as substrate for *Cb4HNR* which reduces it to yield scytalone. Taken together, these results suggest that the *EfPKS1* and *Cppks1* genes that were formerly predicted to be involved with elsinochrome and phleichrome biosynthesis were likely incorrectly annotated in previous publications and are involved in DHN-melanin biosynthesis.

To identify the legitimate elsinochrome and phleichrome PKS genes in *E. fawcettii* and *C. phlei,* respectively, we went back to our KS domain alignment where predicted PKSs CpPTB1 of *C. phlei* and EfETB1 of *E. fawcettii* clustered together with established cercosporin biosynthesis PKSs CTB1 in *C. beticola* and *C. nicotianae,* which hinted at their contribution to perylenequinone biosynthesis (Fig. 3). In line with these initial functional predictions, alignments of the corresponding predicted gene clusters display high similarity and gene conservation within each clade (Fig. 4A). Also, structural differences between perylenequinones can be explained by comparing the predicted metabolite clusters on a gene level. For example, cercosporin and phleichrome only differ in the additional methylenedioxy bridge that is found in the cercosporin molecule (Fig. 1). Accordingly, the predicted phleichrome biosynthesis pathway lacks *CTB9* and *CTB10* that have been shown to be responsible for methylenedioxy bridge formation (de Jonge et al., 2018). Site-directed gene replacement of *EfETB1* in *E. fawcettii* and *CbCTB1* in *C. beticola* led to the successful generation of perylenequinone mutants that are deficient in toxin production under perylenequinone-inducing conditions (Fig. 6A and B). Since SM production relies on different environmental conditions, not every medium is suitable to activate SM production (Calvo et al., 2002; VanderMolen et al., 2013). For *C. beticola,* research on cercosporin-inducing conditions resulted in the identification of thin PDA plates in combination with natural light as the induction condition of choice (Frandsen, 1955; Fajola, 1978; Jenns et al., 1989), which was shown here to stimulate elsinochrome production.

In conclusion, we have shown that it is possible to identify BGCs of structurally related SM compounds based on the phylogenetic relationship of their encompassing PKSs and overall conservation level of the associated cluster genes. By using an established *CTB* gene cluster as reference, it was possible to single out gene clusters responsible for the synthesis of related perylenequinone compounds in different fungal species. Likewise, we successfully identified clusters associated with DHN-melanin production in *C. beticola, E. fawcettii, C. phlei, P. nodorum,* and *S. bambusicola* using the same approach and the confirmed DHN-melanin cluster as input. Future research using this methodology will be useful for the identification of other perylenequinones and their corresponding BGCs in other fungi.

## Experimental procedures

### Elsinoё fawcettii *and* Cladosporium phlei *genome sequencing*

For high-quality genomic DNA extraction of *Elsinoë fawcettii* strain CBS 139.25 and *Cladosporium phlei* strain CBS 358.69, mycelia was scraped from the surface of PDA agar petri dishes and extracted using the CTAB method (Bolton et al., 2016). Library preparation (500 bp) and subsequent paired-end (PE) genome sequencing was done by BGI via the Illumina HiSeq 4000 platform. Approximately 34 million high-quality sequence reads with an average length of 100 bp were generated for both samples, representing 134- and 111-fold coverage for *E. fawcettii* and *C. phlei* respectively. Draft genomes were assembled using SPAdes (version 3.9.0), with default parameters and k-mers 21, 33, 55, 77 and 99. Prediction of protein-coding gene models was performed *ab initio* using the previously prepared *Cercospora beticola* training parameters (de Jonge et al., 2018) in Augustus (version 3.2.1). Genome sequences and annotations are submitted to NCBI and permanently linked on figshare under doi https://doi.org/10.6084/m9.figshare.6173834.

### Secondary metabolite phylogenetic analyses

Phylogenetic analysis of the type I PKS genes and phylogenetic tree analyses were largely performed as described in de Jonge et al. (2018). In short, we used Pfam domain scanning analyses by HMMER3 (Mistry et al., 2013) with hmm profiles for domains PF00109.25 (Beta-ketoacyl synthase, N-terminal domain) and PF02801.21 (Beta-ketoacyl synthase, C-terminal domain) to identify all PKSs in the predicted proteomes of *C. beticola* (09-40), *C. phlei* (CBS 358.69), *E. fawcettii* (CBS 139.25), *S. bambusicola* (Slf14), *P. nodorum* (SN15), *C. heterostrophus* (C5), *A. alternata* (SRC1lrK2f), *A. fumigatus* (Af293), *B. cinerea* (B05.10), Co. *fioriniae* (PJ7), and *M. oryzae* (70-15) that were obtained from NCBI GenBank or Ensemble Fungi. In total we identified 240 proteins across these 11 proteomes. In addition, we added 70 PKSs from Collemare et al. (2014) and Cppks1 (AFP89389.1) from So et al. (2015). All selected proteins for further analyses are listed in Supplementary Table 1. All 311 PKS proteins were subsequently aligned by Mafft (v7.271) using default parameters, after which we extracted the KS domain proportion as previously defined by Pfam scanning. This resulted in an alignment with 311 proteins across 832 positions, that was used to prepare a maximum likelihood phylogenetic tree using RAxML (version 8.2.11), incorporating 100 rapid bootstraps and subsequent automatic, thorough ML search. We then selected the subclass of 94 non-reducing PKSs for further analysis, as defined previously by Kroken et al. (2003). The final phylogenetic tree and figure was prepared in EvolView (Zhang et al., 2012). In this tree, we collapsed the outgroup clade with 20 members containing PKSs involved with citrinin biosynthesis, as indicated in Figure 4. Inclusion in the final set of 74 non-collapsed, non-reducing PKSs is indicated in Supplementary Table 1.

### Secondary metabolite cluster alignment visualization

For comparative analyses of the secondary metabolite clusters across multiple genome sequences we initially identified orthologous protein families across the beforementioned proteomes using orthoFinder (Emms and Kelly, 2015). Subsequently, we used the MultiGeneBlast algorithm (multigeneblast.sourceforge.net), integral part of antiSMASH (Weber et al., 2015), to prepare gene-by-gene cluster alignments across all species and we then re-colored individual genes within each gene cluster according to the protein family analysis.

### Deletion mutant generation

Site-directed gene replacements of *CTB1* and *CbPKS1* in *C. beticola* strain 1-90 and of *EfETB1* and *EfPKS1* in *E. fawcettii* strain CBS 139.25 were generated using the split-marker approach as described in Bolton *et al.,* 2016. Primers are listed in Supl. Table 2. Regardless of phenotype, all putative knockout mutants were screened for site-directed gene replacement. Successful gene deletion was confirmed by the presence of a PCR product using a forward primer upstream of the 5’ flanking region of the target gene design and hygromycin reverse primer MDB-1145. Additionally, absence of an amplicon using target gene-specific primers reconfirmed deletion of the target gene (Supl. Table 2).

### Perylenequinone production assay

Mycelial plugs of 5 mm in diameter from PKS mutant and wild type *C. beticola* and *E. fawcettii* strains were grown on thin potato dextrose agar (PDA, Difco™, BD Diagnostic Systems, Sparks, USA) plates (3.0 mL PDA in a 50 mm Petri plate, amended with 150 μg ml^-1^ hygromycin B (Roche, Mannheim, Germany) (for mutant strains) under a natural light-dark cycle at 21 °C for 7 days. Three 5 mm diameter mycelial plugs of each *E. fawcettii* strain were ground with a micro-pestle in 1 mL potato dextrose broth (PDB, Difco™, BD Diagnostic Systems, Sparks, USA), spread onto thin PDA plates and were grown under 24 hrs light at 28 °C for 7 days.

Total mycelial tissue was excised from the agar plate, blended at high speed for 20 s and extracted with ethyl acetate whilst stirring for 5 min in the dark. Single plate extracts were filtered using two layers of miracloth and dried under a stream of nitrogen (21 °C). The reddish-brown residues were resuspended in 200 μl methanol. Cercosporin concentration was calculated by measuring absorbance at 255 nm using an Agilent Cary 8454 UV-Visible spectrophotometer (Agilent Technologies, Inc., Santa Clara, USA) and 21, 500 as the molar extinction coefficient (Milat and Blein, 1995). Extracts were diluted to ~100 pg μl^-1^ with methanol and centrifuged at 3,000 x g for 5 min. At a minimum, duplicate plate extracts were submitted for mass spectrometric analyses of each fungal strain.

### Mass spectrometric analyses

Positive mode electrospray ionization settings were optimized for cercosporin by infusing a methanolic cercosporin standard (5 ng/μL) (Sigma; St. Louis, USA) into a Waters (Milford, MA) Acquity triple quadrupole mass spectrometer. The precursor ion, product ions, optimum collision energies, and cone voltage were determined by the AutoTune Wizard within the MassLynx 4.1 software (Waters; Milford, MA). Ion transitions used for cercosporin detection were m/z 535 → 415 and m/z 535 → 485 using a cone voltage of 60 and collision energies of 25 and 20 V, respectively.

Elsinochrome standard was not available, therefore an extract from wild type *E. fawcettii* was infused into the mass spectrometer and fragmentation of ions appearing at m/z 547 (the molecular mass of elsinochromes B & D) were optimized using the AutoTune Wizard within the MassLynx 4.1. Presumptive elsinochrome ion transitions used were m/z 547 → 487 and m/z 547 → 457 using a cone voltage of 60 and collision energies of 20 and 35 V, respectively. In some elsinochrome analyses, the mass spectrometer was used as a single sector instrument to collect molecular ions at m/z 547 (elsinochromes B & D), m/z 545 (elsinochrome A), and m/z 549 (elsinochrome C). For both cercosporin and elsinochrome MS/MS experiments, the desolvation temperature was set at 500 °C, and the source temperature was set at 150 °C. Cone gas (N_2_) flow was set at 50 L/h and desolvation gas flow was set at 800 L/h, whereas the collision gas (Ar) flow was 0.16mL/min.

Cercosporin and elsinochrome (isomers B and D) were analyzed using liquid chromatography-tandem mass spectrometry (LC-MS/MS) using a Waters (Milford, MA; USA) Acquity UPLC and Acquity triple-quadrupole mass spectrometer. Data were acquired, processed, and quantified using MassLynx 4.1 with Target Lynx systems. Aliquots of sample extracts (10 μL) were injected onto a 2.1 x 30 mm (1.7 μm) Acquity CSH C18 column protected by a 2.1 x 5 mm CHS guard column (Waters; Milford, MA, USA). Cercosporin and elsinochrome were eluted with a binary gradient consisting of solvent A (0.1% formic acid in pure water) and solvent B (0.1% formic acid in acetonitrile) flowing at 1 mL/min. The gradient program was started at 95% A and transitioned to 25% A over 2 minutes, 5% A at 2.1 minutes, and held at 5% A until 2.5 min when solvent A was ramped back to 95% A at 3 minutes. Solvent composition was held constant until the end of the run time at 4 min. The column temperature was 30°C.

### Melanin production assay

Three mycelial plugs of 5 mm diameter from each of wild type and mutant *C. beticola* strains were ground with a micropestle in 1 mL V8 broth (10% (v/v) clarified V8 juice (Campbell’s Soup Co., Camden, USA), 0.5% (w/v) CaCO_3_) and spread onto single Nylon membranes (Nytran^®^ SuPerCharge Nylon transfer membrane, Schleicher and Schuell, Keene, USA) overlaying V8 agar (as broth but with 1.5% (w/v) agar (BD, Franklin Lakes, USA)) plates (6 mL in a 50 mm Petri plate). *C. beticola* was grown under constant light at 21 °C for 7 days. *E. fawcettii* strains were grown in the same way as *C. beticola* but mycelial plugs were ground in Fries media (as described in Friesen and Faris (2012)) and grown on Fries agar (1.5 % (w/v) agar, 6 mL in a 50 mm Petri plate) at 21 °C under 24 hrs light for 10 days. Total mycelial tissue was excised and weighed before extracting melanin according to Gadd (1982). The tissue was boiled for 5 min in 10 mL distilled water, centrifuged, and the pigment extracted from the supernatant by autoclaving with 3 mL of 1 M NaOH (20 mins, 120 °C). The extract was then acidified to pH 2 with concentrated HCl to precipitate melanin. The precipitate was washed three times with distilled water and dried under a stream of nitrogen (21 °C).

Melanin extracts were solubilized in 2 mL of 2M sodium hydroxide at 50 °C. A spectrophotometric assay was used as described by Kauser et al. (2003) to measure melanin absorbance at 475 nm with a standard curve of synthetic melanin (Sigma-Aldrich, Milwaukee, USA) from 1-100 μg per ml to determine melanin content. The mean melanin content was determined as micrograms of melanin per gram of mycelial tissue for three replicates (individual cultures) and the standard error of the mean calculated. One-way ANOVA was performed with a post-hoc Tukey HSD test to determine differences between the mean melanin contents of wild type strains and each of the three mutants for *C. beticola* and *E. fawcettii,* using a P-value of 0.05 as the significance threshold.

## Acknowledgments

This project was supported in part by USDA-ARS CRIS project 3060-22000-049. Work in the laboratory of B.P.H.J.T. is supported by the Research Council Earth and Life Sciences (ALW) of the Netherlands Organization of Scientific Research (NWO). M.E. was supported by NWO grant 833.13.007. R.d.J. was financially supported by an EMBO Long-Term Fellowship (ALTF 359-2013) and a postdoctoral fellowship of the Research Foundation Flanders (FWO 12B8116N). We thank J. Neubauer (USDA – ARS) for excellent technical assistance. Mention of trade names or commercial products in this publication is solely for the purpose of providing specific information and does not imply recommendation or endorsement by the U.S. Department of Agriculture.

